# mirCCC: Repression-aware graph learning for miRNA-mediated cell-cell communication inference

**DOI:** 10.64898/2026.06.26.734694

**Authors:** Yifan Chen, Jiaming Cui, Shilong Zhang, Enyan Liu, Luyao Xie, Cong Feng, Ming Chen

**Author notes:** (CF); (MC).

## Abstract

Cell–cell communication analyses usually focus on protein ligands and receptors and therefore miss extracellular vesicle transfer of microRNAs, an important route of signalling in cancer. Here we show that microRNA-mediated communication can be inferred from standard single-cell RNA sequencing by detecting coordinated decreases in the expression of validated miRNA target genes. We developed mirCCC, a computational framework that estimates cell-specific microRNA activity, models cellular sending and receiving capacity for extracellular vesicle transfer, and learns
microRNA-resolved communication graphs from transcriptomic data. In synthetic benchmarks with strong confounding signals, mirCCC improved whereas all comparison methods declined. Applied to a human colorectal cancer atlas, mirCCC recovered known colorectal cancer-associated microRNAs and identified stromal- and myeloid-to-epithelial communication converging on a plasticity program linked to TGF-*β* and Wnt/*β*-catenin signalling. These results provide a practical route for studying extracellular vesicle-mediated communication in existing single-cell atlases.

**Author summary:** Cell-cell communication plays important roles across a wide range of biological processes. Most studies of cell-cell communication focus on interactions between ligands and receptors. However, cells can also use extracellular vesicles to deliver microRNAs (miRNAs), which regulate gene activity in receiving cells and contribute to cancer progression, immune regulation, and changes in the tissue environment. Since standard single-cell sequencing does not directly capture miRNAs, this mode of communication remains largely unexplored. Here we developed mirCCC, a computational method for inferring miRNA-mediated communication from standard single-cell transcriptomic data. Instead of detecting miRNAs directly, mirCCC examines the effects they leave in receiving cells. Active miRNAs often cause coordinated suppression of their target genes. mirCCC uses these coordinated reductions in the expression of known target genes to estimate miRNA activity and combines them with the ability of sending cells to package and release miRNAs and of receiving cells to process them. It can therefore reconstruct directional communication networks for individual miRNAs. Our evaluations show that mirCCC remains reliable in the presence of misleading signals and reveals biologically meaningful regulatory patterns in human colorectal cancer data. Overall, mirCCC provides a practical way to study miRNA-mediated cell-cell communication using existing single-cell transcriptomics datasets.

## Introduction

Cell-cell communication (CCC) analysis has developed into a rich computational ecosystem. CellChat combines ligand–receptor prior knowledge with a mass-action model to analyze communication probabilities and network patterns [1]; NicheNet links sender-side ligand signals to receiver-side transcriptional responses [2]. LIANA+ integrates multiple CCC resources into a unified framework [3]; 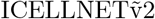 emphasizes multi-subunit ligand/receptor complexes [4]. More recent methods extend cell–cell communication analysis beyond classical ligand–receptor models. DeepTalk extends CCC to spatial transcriptomics by integrating spatial context into graph-based inference [5]. CellNEST models relay communication to capture multi-step signaling cascades across intermediate cell states [6]. SigXTALK further expands CCC analysis to pathway-level crosstalk by linking signaling pathways across interacting cell populations [7]. Despite these advances, most of these methods still rely on protein ligand–receptor interactions as the underlying paradigm.

However, cell-cell communication is not limited to protein ligand–receptor signaling. Extracellular vesicles (EVs) serve as long-range carriers transferring miRNAs, proteins, and lipids between cells [8], playing widespread roles in immune evasion, metastasis, microenvironment remodeling, and therapy resistance [9]. EV-mediated miRNA transfer is functionally distinct from ligand–receptor signaling because it acts through post-transcriptional gene repression rather than receptor activation, making it difficult to characterize using existing CCC frameworks [10]. Furthermore, standard single-cell transcriptomic workflows rely on poly(A)-based capture and cannot directly measure mature miRNAs [11].

Computational tools for inferring EV-derived miRNA communication at the single-cell level remain scarce. miRTalk was the first to formalize this task, integrating scRNA-seq data with EV-related prior information and miRNA-target interaction databases [12]. However, miRTalk relies on detected miRNA expression in sender cells, whereas conventional poly(A)-based scRNA-seq provides limited coverage of mature miRNAs. MIR-gene transcripts may not faithfully reflect mature miRNA activity or EV loading. In addition, miRTalk mainly summarizes miRNA–target interactions between annotated cell types, limiting its ability to resolve cell-to-cell heterogeneity and reconstruct miRNA-specific communication links at single-cell resolution.

Based on these considerations, we propose mirCCC, a repression-aware graph learning framework for inferring miRNA-mediated cell–cell communication from standard scRNA-seq data. Rather than treating miRNA communication as another class of ligand–receptor events, mirCCC infers communication from receiver-side target-gene repression patterns consistent with miRNA activity. It first employs signed diffusion on a miRNA–target bipartite graph to estimate cell-level miRNA activity proxies from target-gene expression, thereby avoiding reliance on direct miRNA detection in conventional poly(A)-based scRNA-seq. It then estimates cellular sending and receiving capacities from the expression of EV biogenesis, miRNA sorting, and RISC pathway genes. On this basis, mirCCC constructs directed graphs for individual miRNAs to resolve communication heterogeneity among cells within the same annotated cell type and to learn representations of their miRNA-mediated communication context. Crucially, communication strength is determined by miRNA-specific proxy–repression matching: a high sender-side proxy for a given miRNA must coincide with repression of its corresponding target genes in the receiver cell. This requirement reduces false positives caused by co-expression, cell-state covariation, or nonspecific activation of EV-related machinery. Overall, mirCCC extends single-cell communication inference beyond protein ligand–receptor signaling and enables the analysis of miRNA-mediated regulation using routine scRNA-seq data.

## Methods

An overview of the full pipeline is shown in Fig. 1.

**Fig 1.**
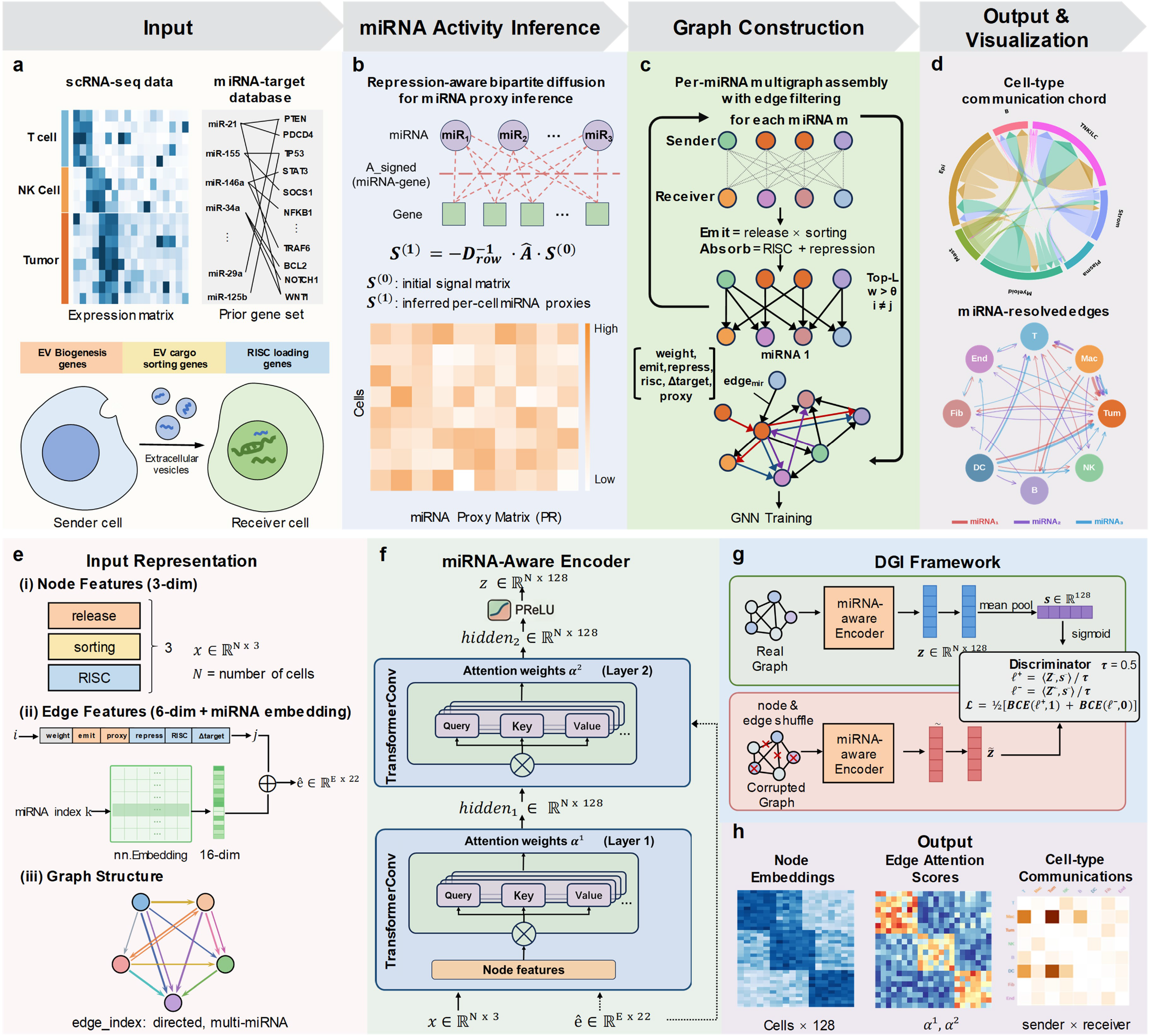
mirCCC pipeline overview and model architecture. **(a)** Input data: scRNA-seq expression matrix and curated miRNA–target interaction database, together with three prior gene sets (EV biogenesis, EV cargo sorting, and RISC loading genes) that define the molecular machinery for EV-miRNA communication. **(b)** miRNA activity inference: a repression-aware signed bipartite diffusion propagates z-scored target-gene expression through a weighted miRNA–gene adjacency matrix to produce per-cell miRNA proxy scores. **(c)** Per-miRNA directed graph construction: for each candidate miRNA, sender and receiver cells are selected based on emission and absorption capacity, respectively, and connected by directed edges carrying 6-dimensional continuous features plus a learnable miRNA embedding. **(d)** Output and visualization: cell-type-level communication matrices and miRNA-resolved chord diagrams. **(e)** Input representation for the graph neural network: 3-dimensional node features (release, sorting, RISC scores), 22-dimensional augmented edge features (6 continuous + 16-dim miRNA embedding), and directed multi-miRNA graph structure. **(f)** miRNA-aware encoder architecture: two stacked TransformerConv layers with multi-head attention incorporating edge features, producing 128-dimensional node embeddings. Attention weights *α*^1^ and *α*^2^ from both layers are retained for downstream scoring. **(g)** Self-supervised training via Deep Graph Infomax (DGI): a cosine-similarity discriminator contrasts node embeddings from the real graph against those from a corrupted graph (generated by independent node and edge feature shuffling) to learn meaningful representations without labeled data. **(h)** Model outputs: node embeddings, edge-level attention scores, and aggregated cell-type communication matrices.

### Prior knowledge databases

mirCCC relies on four curated gene sets as prior knowledge inputs (Fig. 1a). Experimentally validated miRNA-target interactions for extracellular vesicle (EV)-derived miRNAs were obtained from miRTalkDB, the curated database underlying miRTalk [12]. miRTalkDB integrates experimentally verified EV-derived miRNAs curated from EV-focused resources, including EVmiRNA [13], ExoCarta [14], and Vesiclepedia [15], together with experimentally validated miRNA-target interactions from miRTarBase [16] and TarBase [17]. miRTalk further assigns confidence annotations to these interactions, including Functional MTI, Functional MTI (Weak), and Non-Functional MTI. Three additional pathway gene lists were used to define the EV-miRNA communication machinery: EV biogenesis, miRNA sorting, and RISC-mediated target silencing. The complete gene lists are distributed with the pipeline repository. All gene names are unified to HGNC symbols via an alias-mapping procedure prior to analysis.

### miRNA activity inference via signed graph diffusion

Standard scRNA-seq protocols rely on polyA-tail capture and thus cannot directly measure mature miRNAs, which are approximately 22 nucleotides long and lack polyA tails. Since active miRNAs leave a detectable repression footprint on their target genes, we reverse-infer cell-level miRNA activity directly from mRNA expression data (Fig. 1b).

We construct a weighted bipartite graph *G* = (*M ∪T, E*) from miRTalk, where *M* is the set of miRNAs and *T* is the set of target genes present in the expression data. Interactions are filtered by species (human) and minimum target count (*≥* 10 per miRNA). Each edge (*m, t*) *∈E* is assigned a confidence weight *w*_*mt*_ reflecting the strength of experimental evidence:

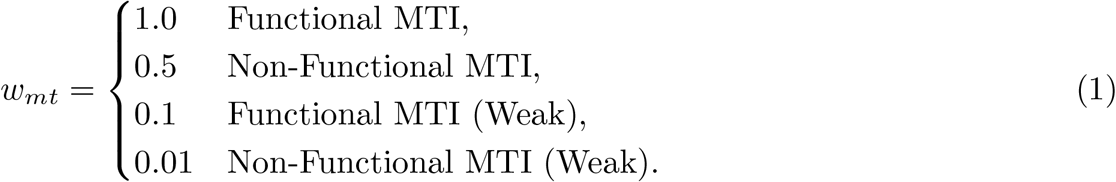

These raw weights are assembled into a biadjacency matrix **A**_raw_ ∈ℝ^|*M* |*×*|*T* |^. To mitigate the influence of promiscuous target genes regulated by many miRNAs, we apply inverse document frequency (IDF) weighting followed by L1 row-normalization. IDF down-scales columns corresponding to genes targeted by many miRNAs:

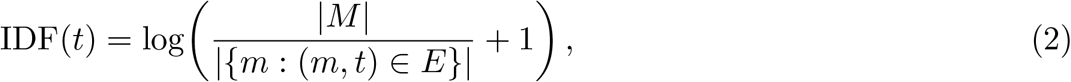

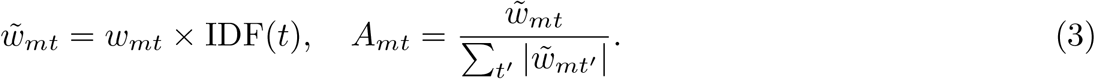

Let 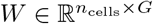 denote the expression matrix restricted to target genes retained after preprocessing. For each target gene, expression values are standardized across cells to obtain a z-scored matrix *W*_*z*_. These gene-level signals are then placed on the gene side of the miRNA–target bipartite graph, forming the initial signal matrix

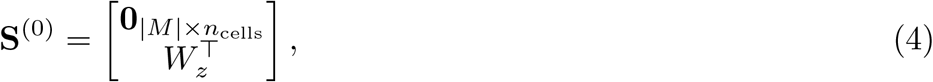

where the miRNA block is initialized to zero and the gene block contains the transposed target-gene z-scores.

Let **A**_signed_ *∈* ℝ^|*M* |*×G*^ denote the signed, weighted miRNA–target adjacency matrix. To propagate repression signals from target genes to miRNAs, we construct the full bipartite adjacency

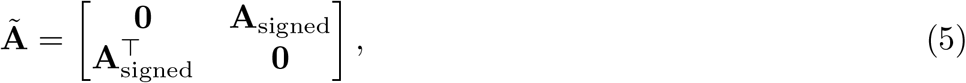

and apply symmetric degree normalization,

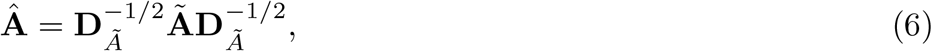

where **D**_*Ã*_ is the diagonal degree matrix of **Ã**. A single-step signed diffusion is then performed as

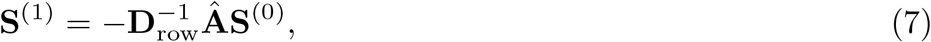

where **D**_row_ = diag(|**Â**| **1**) is a row-wise normalization term. This additional normalization improves numerical stability and reduces bias arising from variation in miRNA target-set size.

The negative sign encodes the repressive prior of miRNA regulation: an active miRNA is expected to be associated with reduced expression of its target genes, such that lower target-gene z-scores are transformed into higher inferred miRNA proxy values after propagation. The miRNA block of **S**^(1)^ is then extracted and arranged into the proxy matrix

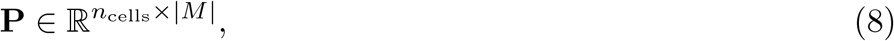

which stores the inferred activity proxy of each miRNA in each cell.

To assess target observability, we compute a coverage quality score for each miRNA based on the fraction of support-weighted target evidence retained in the expression matrix. Let *S*_all_(*m*) denote the total support weight of all annotated targets of miRNA *m*, and let *S*_obs_(*m*) denote the total support weight of those targets with detection fraction at least 0.05 across cells. We define

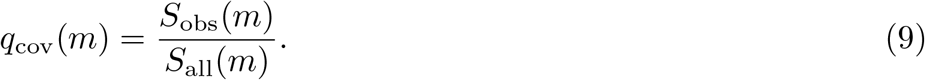

miRNAs with low target coverage are then down-weighted by multiplying their diffused scores by *q*_cov_(*m*)^*γ*^, where *γ* = 2.5 by default. In downstream graph construction, each miRNA-specific proxy vector is further min–max normalized across cells to [0, 1], yielding proxy_01_(*m, c*).

### Sender and receiver capacity scoring

To quantify sender- and receiver-side molecular capacity, we computed pathway activity scores for the EV biogenesis, miRNA sorting, and RISC gene sets using mean z-score aggregation of log-normalized expression. Raw counts were library-size normalized to 10^4^ counts per cell and log-transformed using Scanpy [18]. For each gene *g*, expression was standardized across cells as

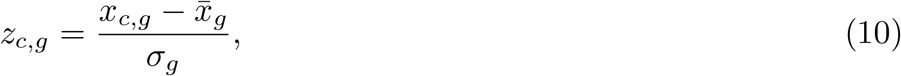

where 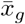 and *σ*_*g*_ denote the mean and standard deviation of gene *g* across all cells. For each pathway gene set *G*_*k*_, the per-cell pathway score was defined as

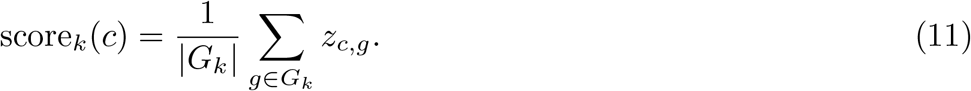

This yielded three pathway activity scores per cell: a release score from EV biogenesis genes, a sorting score from miRNA sorting genes, and a RISC score from RISC pathway genes. Each score was subsequently min–max normalized to [0, 1], yielding release_01_(*c*), sorting_01_(*c*), and risc_01_(*c*).

To define sender-side EV-miRNA emission capacity, we required that a cell exhibit both EV biogenesis activity and miRNA sorting activity. Accordingly, the sender emission prior was defined as

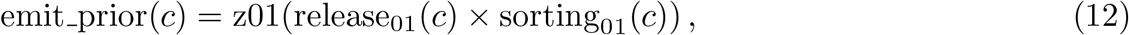

where z01(·) denotes min–max normalization to [0, 1] across cells. Intuitively, this score assigns high values only to cells that jointly exhibit strong vesicle production and active miRNA loading machinery. In contrast, risc_01_(*c*) was used as the receiver-side machinery score, reflecting the cellular capacity to process incoming miRNAs and mediate downstream target repression.

### Per-miRNA directed graph construction

A key design principle of mirCCC is that miRNAs are modeled independently rather than collapsed into a single aggregate communication score (Fig. 1c). For each candidate miRNA, we constructed a miRNA-specific directed sender-to-receiver edge set over the shared cell population and retained the miRNA identity of each edge in the pooled graph representation. For each miRNA, sender cells were prioritized by combining the cell-level emission prior with the normalized miRNA proxy, whereas receiver cells were prioritized by the inferred target-repression score. We then selected the top sender and receiver candidates and, for each sender, retained only the top receiver partners, yielding a sparse directed graph focused on the most plausible communication routes. Self-loops and weakly supported edges were removed. Each retained edge was annotated with six features describing matching strength, sender capacity, receiver state, and miRNA-specific activity, whereas each node carried three pathway-based features corresponding to EV biogenesis, miRNA sorting, and RISC activity.

#### Node features

Each cell was represented by three pathway-based node features: normalized EV biogenesis activity, normalized miRNA sorting activity, and normalized RISC pathway activity (Fig. 1e).

#### Sender selection

For each miRNA *m*_*k*_, a miRNA-specific sender score was computed for every cell as the product of the cell-level emission prior and the normalized proxy of *m*_*k*_:

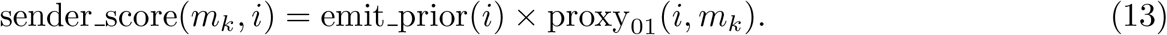

This favors cells that both exhibit EV-miRNA emission capacity and show evidence of activity for the corresponding miRNA. The top-*S* cells were then selected as sender candidates, where *S* is a configurable hyperparameter.

#### Receiver scoring

Receiver candidates were prioritized using a hybrid repression score that combines two complementary measures of target-gene suppression through their geometric mean:

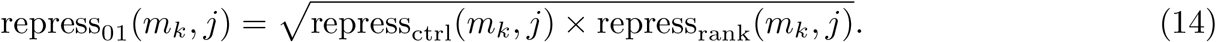

The first component was a control-adjusted target-module score. For each miRNA, the weighted mean expression of its target genes in a given cell was compared with that of expression-matched control genes selected from the same global expression bins, yielding an adjusted target-module signal. Lower adjusted target-module expression corresponded to stronger inferred repression and was rescaled to [0, 1] to obtain repress_ctrl_.

The second component was a within-cell rank-based repression score computed from the weighted normalized ranks of target genes. This term captures ordinal repression patterns and is less sensitive to outliers in absolute expression magnitude. The geometric mean of the two components therefore combines absolute repression relative to matched controls with rank-based evidence of target suppression.

To incorporate receiver-side processing capacity, the final receiver absorption score was defined as

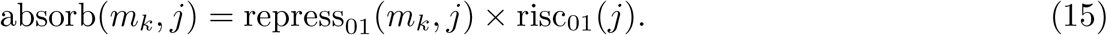

The top-*R* cells were then selected as receiver candidates, where *R* is a configurable hyperparameter (*R* = 150 in the default configuration). For each sender cell, directed edges were retained only to its top-*L* receiver candidates, where *L* is also configurable (*L* = 10 by default). Candidate edges with low combined sender–receiver weight were discarded using a minimum threshold of 0.005.

#### Edge features

Each directed edge (*I → j*) associated with miRNA *m*_*k*_ carried a six-dimensional continuous feature vector comprising the base sender–receiver matching weight, the sender’s emission prior, the receiver’s repression score, the receiver’s RISC capacity, the receiver-side control-adjusted target-module signal, and the sender’s normalized miRNA proxy score. In addition, each edge carried a discrete miRNA index *k*, which was used for learnable miRNA embedding lookup in the graph encoder.

### miRNA-aware Graph Transformer encoder

The miRNA-aware encoder jointly integrates cell-level node features with miRNA-specific edge information (Fig. 1f). Each miRNA index is first mapped to a learnable embedding of dimension 16, which is concatenated with the 6-dimensional continuous edge feature vector to form an augmented edge representation.

The encoder consists of two stacked `TransformerConv` layers [19], each using multi-head attention with edge features incorporated into the attention computation. In the default configuration, we used 4 attention heads and a hidden dimension of 128, with multi-head outputs aggregated rather than concatenated. A PReLU activation was applied after the second layer to obtain the final node embeddings **z**_*i*_ *∈* ℝ^128^.

Attention weights from both layers were retained for downstream analysis, with final-layer attention used in communication score extraction.

### Self-supervised training via Deep Graph Infomax

Because ground-truth labels for miRNA-mediated communication are unavailable, we trained the encoder using Deep Graph Infomax (DGI) [20], a self-supervised objective that maximizes agreement between local node representations and a global graph summary (Fig. 1g). Following the standard DGI formulation, the graph-level summary vector was computed by applying a sigmoid activation to the mean of all node embeddings.

Negative examples were generated by a corruption function that independently permuted node features and continuous edge attributes while keeping the graph topology fixed. This dual corruption yields a stronger contrastive signal than perturbing only one modality, as the encoder must learn the correspondence between cellular states and their miRNA-aware communication context. Training then followed a contrastive objective that distinguishes node embeddings from the original graph from those obtained under corruption. Specifically, embeddings and the graph summary vector were L2-normalized, and a cosine-similarity discriminator with temperature *τ* = 0.5 was used:

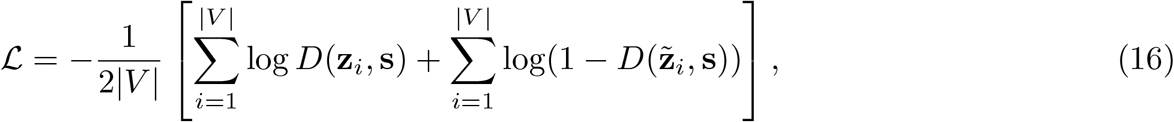

where 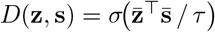 with 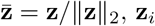, **z**_*i*_ and 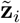 denote node embeddings from the original and corrupted graphs, respectively, and **s** = *σ*(mean_*i*_ **z**_*i*_) denotes the graph summary vector.

#### Optimization

The model was optimized with Adam using an initial learning rate of 0.001 and a cosine annealing schedule with a minimum learning rate of 0.01 *× η*_0_. Mini-batch training was performed with NeighborLoader using two-hop neighborhood sampling with 10 neighbors per hop and a batch size of 1024, with receiver nodes used as input seeds. The number of training epochs was treated as a configurable hyperparameter.

### Communication score computation

After training, miRNA-specific communication scores were computed for directed sender–receiver edges by combining hand-crafted biological priors with learned attention-based modulation (Fig. 1h). For a directed edge (*i → j*) associated with miRNA *m*_*k*_, the final score was defined as

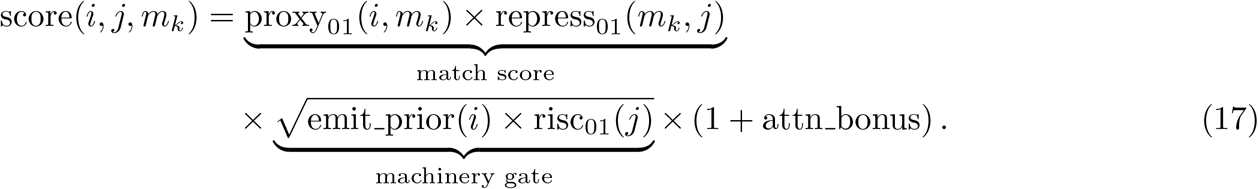

The match score captures miRNA-specific sender–receiver compatibility by combining the sender-side proxy of *m*_*k*_ with the receiver-side repression signal for the same miRNA. The machinery gate acts as a soft biological filter, increasing scores only when the sender exhibits EV-miRNA emission capacity and the receiver exhibits RISC loading capacity. Learned attention then provides a mild multiplicative refinement:

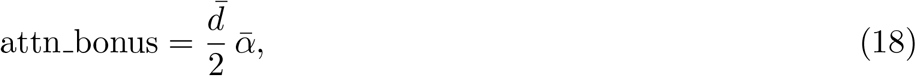

where 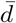 denotes the average graph degree and 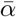 denotes the mean edge attention derived from the retained transformer attention weights. The final communication score therefore combines miRNA-specific matching, sender–receiver molecular competency, and graph-learned contextual refinement.

#### Aggregation

For downstream summarization, miRNA-specific edge scores were first aggregated at the cell-pair level by combining all miRNA-mediated edges between the same sender and receiver cells. In the default setting, this aggregation was performed by summation. Cell-type-level communication matrices were then obtained by summing the resulting cell-pair scores over all constituent sender–receiver cell pairs belonging to cell types *A* and *B*, respectively (Fig. 1d,h).

### Synthetic benchmark data

To evaluate mirCCC under controlled conditions, we generated two synthetic scRNA-seq benchmark settings, each containing 2,000 cells, 8 cell types, 2,000 genes, and 3 implanted communication patterns. In the Base setting, each communication pattern was mediated by 16 ground-truth miRNAs. In the Trap setting, each communication pattern was mediated by 12 ground-truth miRNAs. Communication signals were injected by increasing EV biogenesis and miRNA sorting activity in sender cells, increasing RISC activity in receiver cells, and repressing the corresponding miRNA target genes in receivers. Dropout and expression noise were added to model technical variability.

Two benchmark settings were used. The *Base* setting featured uniform miRNA importance, no orthogonal target-selection trap, no per-miRNA gradient, and predominantly easy negatives (90%). The *Trap* setting introduced orthogonal target selection, variable miRNA importance, and graded per-miRNA signals, with easy negatives reduced to 45%, increasing hard cross-pattern confounders.

Six methods were evaluated: mirCCC and five baselines, including two simple heuristics (Prior-Product, Matched-Product), one miRNA-oriented (miRTalk-Restored), and two adapted ligand–receptor-style methods (CellChat-style, NicheNet-style). All were run under the same benchmark framework. miRTalk-Restored is a paper-faithful reimplementation restoring EVBS-based sender scoring, RISC/RITAC-based receiver scoring, rank-based target evaluation, optional permutation testing, and specificity calculation, while relying on proxy miRNA activity. The CellChat-style and NicheNet-style baselines are miRNA-adapted analogs of classical ligand–receptor methods. Missing ground-truth pairs were completed using the method-specific fill score defined below, and results were aggregated across user-specified random seeds and settings.

### Benchmark methods and unified evaluation

To ensure comparability across methods, all predictions were converted into a unified cell-pair-level format containing the sender cell, receiver cell, and communication score. The resulting predictions were merged with the ground-truth cell-pair labels used in the synthetic benchmark. Ground-truth pairs not covered by a given method were assigned a score no greater than the minimum score produced by that method. Performance was subsequently evaluated at both the cell-pair and cell-type levels.

In addition to mirCCC, we implemented five comparison methods. Matched Product calculates the product of a sender-cell EV-miRNA emission score and the mean receiver-side repression score derived separately for each miRNA-specific target set. Prior Product instead multiplies the sender-side prior score by a global receiver-side repression score calculated from the union of all candidate miRNA target genes. Neither method includes graph representation learning. The distinction between them is that Matched Product preserves the correspondence between individual miRNAs and their respective target sets before aggregation, whereas Prior Product collapses all target genes into a single global receiver-side score.

miRTalk-Restored is a benchmark-oriented reimplementation based on the methodological description of miRTalk [12], rather than a direct execution of the official software. The original miRTalk workflow relies on detectable sender-side miRNA expression and primarily summarizes communication at the cell-type level, whereas our benchmark uses standard scRNA-seq data without direct mature-miRNA measurements and requires scores for predefined individual sender–receiver cell pairs. We therefore reimplemented its core scoring principles within the unified benchmark framework. The adapted implementation retains the principal components of miRTalk, including EV biogenesis and sorting-related sender modules, RISC/RITAC-related receiver modules, miRNA proxy scores, and receiver scoring based on target-gene expression ranks. For each candidate cell pair, miRNA-specific communication scores were calculated, and the maximum score across candidate miRNAs was used as the final cell-pair prediction.

CellChat-style and NicheNet-style were constructed as conceptual adaptations of CellChat [1] and NicheNet [2], respectively, for the miRNA communication setting. In CellChat-style, sender-side miRNA emission and receiver-side target repression replace conventional ligand and receptor signals, and communication scores are calculated using a mass-action formulation with a Hill function at the cell-type level. In NicheNet-style, a miRNA-target regulatory-potential matrix is constructed and combined with receiver-side target-gene repression patterns to estimate miRNA regulatory activity, which is subsequently multiplied by the sender-side emission score to obtain a cell-pair communication score.

The official CellChat and NicheNet implementations were not directly applied because their original models are designed for protein ligand-receptor communication and do not directly accept miRNA proxies, miRNA-target repression evidence, or EV-related sender and receiver features as inputs. Similarly, the original miRTalk workflow relies on detectable sender-side miRNA expression and primarily summarizes communication at the cell-type level, whereas the present benchmark uses standard scRNA-seq data without direct mature-miRNA measurements and defines ground truth at the individual cell-pair level. We therefore implemented task-specific adaptations based on the core modeling principles of each method and standardized their candidate cell pairs, output structure, and evaluation procedure. Full scoring definitions, parameter settings, output-completion procedures, and implementation details for all benchmark methods are provided in S1 Appendix and the accompanying code repository.

#### Enrichment analysis

To assess whether the top-ranked miRNAs were supported by prior knowledge, we tested the enrichment of literature-curated CRC-associated miRNAs among the top candidates. Specifically, the top 20% of miRNAs ranked by mirCCC score along each sender–receiver axis were compared to the full set of candidate miRNAs using Fisher’s exact test to evaluate whether literature miRNAs were overrepresented. P-values were reported for each axis to quantify the significance of enrichment.

### Evaluation metrics

Model performance was primarily evaluated using AUROC (area under the receiver operating characteristic curve) and AUPRC (area under the precision–recall curve) for binary classification of ground-truth communication cell pairs in the synthetic benchmark. For fair comparison, all methods were evaluated on the same labeled cell-pair set after completing missing predictions using the method-specific fill score defined above.

### Implementation

mirCCC is implemented in Python using PyTorch and PyTorch Geometric for model training, scanpy and anndata for scRNA-seq data handling, and scipy for sparse matrix operations. All experiments were conducted on a single NVIDIA GPU. The source code, installation instructions, and example workflows are available at https://github.com/yfchen801/mirCCC.

## Results

### Synthetic benchmarking shows robustness to confounding signals

To systematically evaluate the inference accuracy and robustness of mirCCC, we benchmarked it under two synthetic data settings (see Methods). Both settings contained three ground-truth miRNA communication axes. In the Base setting, each axis was mediated by 16 miRNAs selected from the miRTalk resource [12]; sender cells showed increased EV biogenesis and miRNA sorting activity, whereas receiver cells showed increased RISC activity and systematic repression of the corresponding miRNA target genes. In the Trap setting, each axis was mediated by 12 miRNAs and included three additional sources of difficulty: (i) orthogonal target selection reduced target-gene overlap across communication axes, thereby requiring miRNA-specific resolution; (ii) variable importance assigned stronger repression signals to 35% of the implanted miRNAs and weaker signals to the remainder; and (iii) the proportion of hard negative cell pairs increased from 10% to 55%, such that more negative pairs exhibited sender- or receiver-associated machinery signals without belonging to a true communication pair. Each setting was generated using three independent random seeds, yielding six experiments in total.

We compared mirCCC against five methods: miRTalk [12], the only existing miRNA communication inference tool, along with CellChat-style [1], NicheNet-style [2], matched product, and prior product baselines. To ensure fair comparison, all methods were evaluated on identical ground-truth cell pairs; for pairs not covered by a given method, predictions were completed using the method-specific fill score defined in Methods.

Under the Base setting, mirCCC achieved an AUROC of 0.884*±* 0.014 (AUPRC 0.892 *±*0.010; mean ± SD over three runs), which was lower than other methods (AUROC 0.911–0.951), indicating relatively conservative scoring in the absence of confounders (Fig. 2a–b). In contrast, under the Trap setting, mirCCC was the only method whose performance improved, with an AUROC of 0.926 *±*0.020 (absolute increase Δ = +0.048), whereas all other methods declined relative to Base: miRTalk by 0.056, CellChat by 0.209, NicheNet by 0.215, Matched Product by 0.238, and Prior Product by 0.240 (Fig. 2c–f).

**Fig 2.**
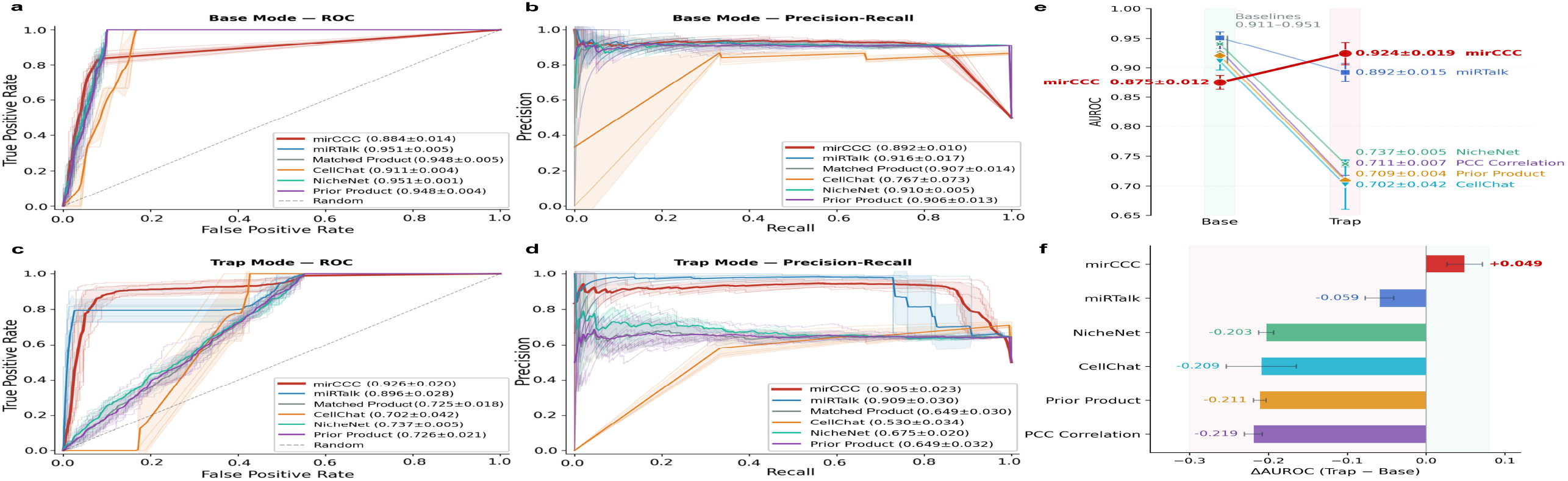
Cell-pair-level synthetic benchmark performance under standard and confounded settings. (a,b) Receiver operating characteristic (ROC) and precision–recall curves in the Base setting. (c,d) ROC and precision–recall curves in the Trap setting. Base denotes the standard synthetic benchmark, whereas Trap denotes the confounded benchmark with deliberately misleading non-specific sender/receiver signals. Methods are color-coded as mirCCC (red), miRTalk (blue), Matched Product (grey), CellChat (orange), NicheNet (teal), and Prior Product (purple). In (a–d), darker lines indicate mean curves across runs and lighter lines indicate individual runs; dashed diagonals indicate random classification. (e) Summary of area under the ROC curve (AUROC) in the Base and Trap settings; points indicate mean values and error bars indicate standard deviation across runs. (f) Change in AUROC between settings (ΔAUROC = Trap *−* Base); bars indicate mean change and error bars indicate standard deviation. AUPRC, area under the precision–recall curve.

This divergence reflects a fundamental difference in method design. The baseline methods (CellChat, NicheNet, Matched Product, Prior Product) rely on sender–receiver expression correlations or machinery gene expression levels; decoy cells satisfy these surface-level conditions, leading to false-positive communication calls. In contrast, mirCCC’s repression-aware scoring infers communication by detecting systematic suppression of target genes in receiver cells. Although decoy cells express the requisite machinery genes, they lack genuine miRNA–target repression signals and therefore do not yield high scores. These results demonstrate that mirCCC’s repression-aware scoring provides robustness against confounding signals that mislead existing approaches in the Trap scenario.

### CRC communication landscape reveals an Epi-centered receiver context

To evaluate mirCCC on real data, we applied it to the Pelka et al. CRC single-cell atlas (GSE178341) [21], selecting MMRp patient C162 comprising 9,994 cells across 7 major cell lineages: Epi, Myeloid, Strom, T/NK/ILC, B, Plasma, and Mast (Fig. 3a). miRNA–target interactions were obtained from miRTalk [12], based on experimentally validated data from miRTarBase [16], retaining only 12,731 Functional MTI-level records. After coverage filtering, 185 miRNAs entered the analysis.

**Fig 3.**
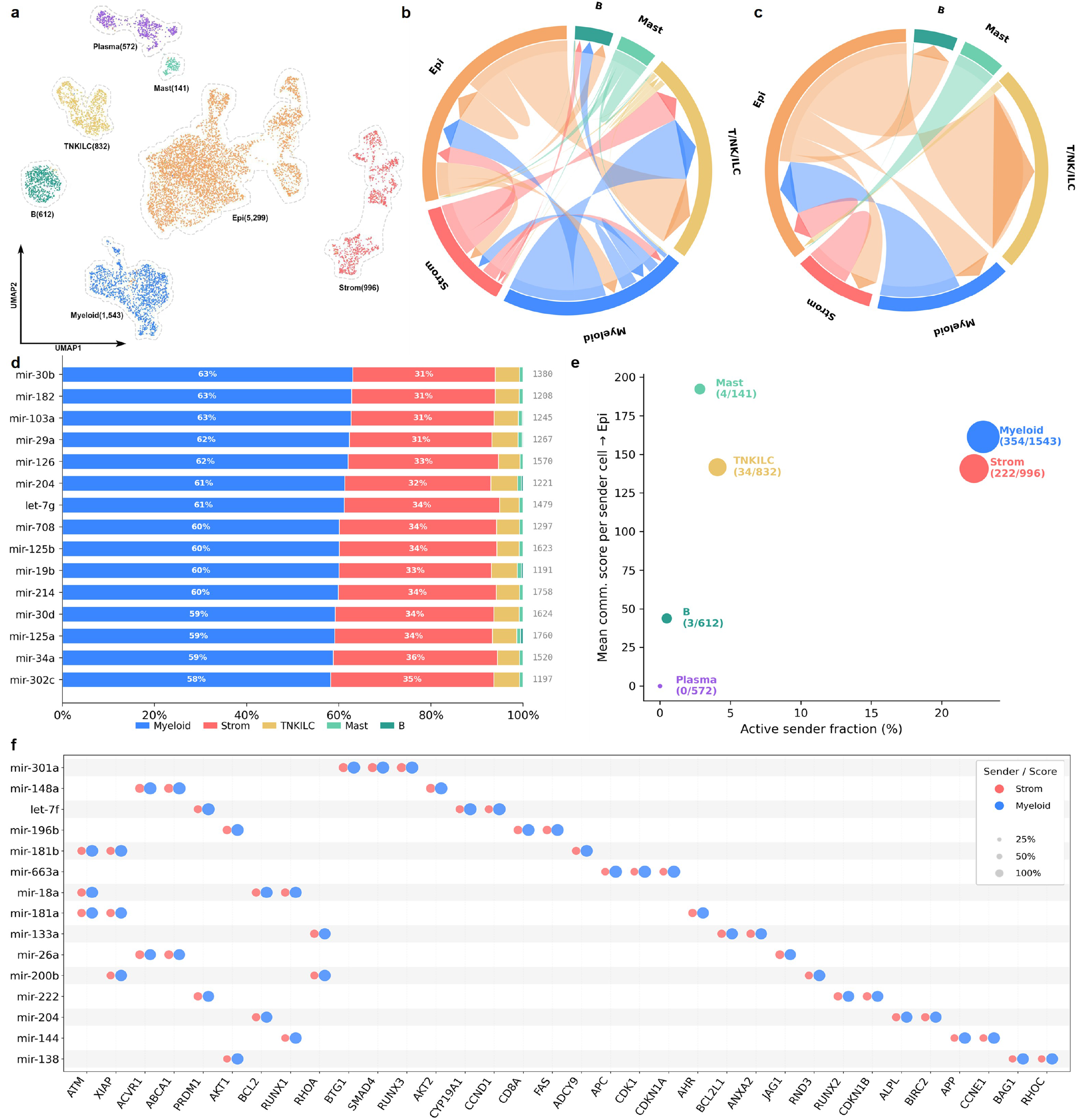
CRC communication landscape inferred by mirCCC in patient C162. **(a)** UMAP visualization of 7 major cell lineages; cell counts are shown in parentheses. **(b)** Global miRNA-mediated cell-cell communication chord diagram; arc length is proportional to communication flow and arrowheads indicate directionality. **(c)** Epi-centric paracrine chord diagram, showing the communication contribution from each non-Epi cell type toward Epi. **(d)** Stacked percentage bar chart of sender cell type contribution per miRNA for paracrine communication toward Epi; numbers on the right indicate absolute communication scores. **(e)** Paracrine sender capacity scatter plot for non-Epi cell types; *x*-axis: fraction of cells selected as active senders, *y*-axis: mean communication score per sender cell toward Epi, bubble size proportional to the number of sender cells. **(f)** Split bubble plot showing the 15 miRNAs with the highest combined communication scores across the Strom *→*Epi and Myeloid *→*Epi axes and their associated target genes. Bubble size represents the sender-specific communication score, and color indicates the sender compartment (red, Strom; blue, Myeloid).

The global communication network inferred by mirCCC revealed that Epi cells serve as a major receiver of miRNA signals, simultaneously accepting inputs from multiple non-epithelial compartments (Fig. 3b). An Epi-centric paracrine communication view further delineated the relative contributions of each sender (Fig. 3c). Quantification of sender-specific contributions showed that Myeloid and Strom together accounted for approximately 94% of the total paracrine communication score directed toward Epi (Myeloid 60.7%, Strom 33.2%), far exceeding T/NK/ILC (5.1%), Mast (0.8%), and B (0.1%) (Fig. 3d). This pattern was remarkably consistent across all top 15 miRNAs, with each showing a Myeloid-plus-Strom contribution of 90–95%. The dominance of these two compartments was further supported by a two-dimensional analysis of sender capacity. Myeloid and Strom had the highest fractions of active sender cells (23.0% and 22.3%, respectively) and exhibited the highest mean communication score per sender cell directed toward Epi (Fig. 3e).

Despite the quantitative difference between Myeloid and Strom in overall sender contribution, the miRNA–target circuit diagram revealed a key qualitative feature: the top-ranked miRNAs on the Strom *→* Epi and Myeloid *→* Epi axes were highly overlapping (Fig. 3f). The 15 miRNAs with the highest combined scores generally received substantial contributions from both stromal and myeloid senders. This cross-compartment convergence suggests that stromal and myeloid cells may deliver regulatory signals to epithelial cells through a shared miRNA repertoire. Based on these observations, subsequent analyses focus on the Strom*→*Epi and Myeloid*→*Epi axes to systematically assess the biological plausibility of mirCCC predictions.

### Convergent stromal and myeloid miRNAs target an epithelial plasticity module

To evaluate whether mirCCC-prioritized candidate miRNAs along the Strom *→* Epi and Myeloid *→* Epi axes are supported by prior CRC biology, we tested their enrichment among literature-curated CRC-associated miRNAs. On both axes, these literature-curated CRC-associated miRNAs were 3.2-fold enriched within the top 20% of mirCCC-ranked miRNAs (7/11, Fisher’s exact *p* = 1.1 *×* 10^*−*3^; Fig. 4b). A similar enrichment was observed on the Epi *→* TNKILC axis (7/15, 2.3-fold, *p* = 0.013), whereas the Myeloid *→* TNKILC axis showed no significant enrichment, serving as an internal negative control. These results indicate that mirCCC recovers biologically relevant candidates without explicitly incorporating prior knowledge of miRNA function during inference.

**Fig 4.**
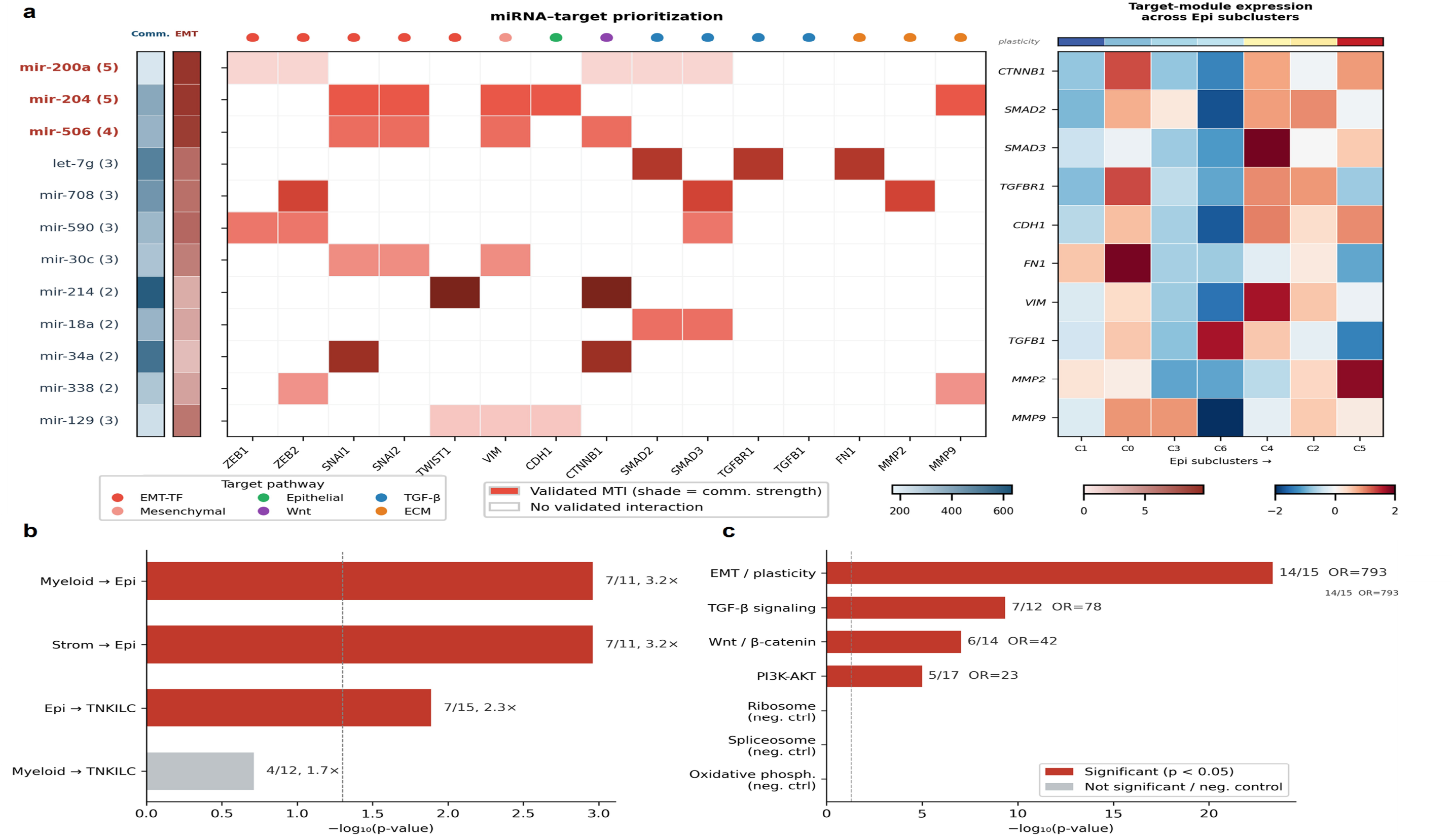
Functional characterization of stromal- and myeloid-derived miRNAs converging on epithelial plasticity. (a) miRNA-target prioritization matrix with epithelial subcluster expression context. (b) Enrichment of literature-curated CRC-associated miRNAs among top-ranked candidates across axes. (c) Pathway enrichment specificity and negative-control pathway comparison for predicted targets.

More importantly, the predicted targets of the prioritized miRNAs were not dispersed across unrelated genes but preferentially converged on an epithelial plasticity-related regulatory module. The miRNA–target prioritization matrix (Fig. 4a) displays 12 representative candidates, comprising three miRNAs highlighted for their broad coverage of the curated plasticity gene set–miR-200a, miR-204, and miR-506–and nine additional high-ranking candidates. Specifically, miR-200a targets ZEB1, ZEB2, SMAD2, SMAD3, and CTNNB1; miR-204 targets SNAI1, SNAI2, VIM, CDH1, and MMP9; and miR-506 targets CTNNB1, SNAI1, SNAI2, and VIM. Collectively, these three miRNAs cover 14 of the 15 curated EMT/plasticity-related genes, forming a coherent target module centered on epithelial plasticity regulation. These predictions are broadly consistent with prior experimental evidence: miR-200a is a canonical EMT suppressor acting through the miR-200/ZEB double-negative feedback loop. Additional literature support among the displayed candidates includes the involvement of miR-34a in EMT-associated invasion programs in CRC and experimentally validated transfer of CAF-derived exosomal miR-200b-3p, supporting the relevance of the miR-200 family in this context. Among the 12 displayed candidates, eight have direct or family-level literature support in CRC or EMT contexts (Table 1). The complete rankings for these two epithelial-directed axes, together with the Epi-to-TNKILC and Myeloid-to-TNKILC comparison axes, are provided in S1 Table. The table includes total communication scores, edge counts, mean scores per edge, validated target counts, and EMT/plasticity-related target annotations.

**Table 1.** Literature support for selected top-ranked miRNAs on the Strom *→*Epi and Myeloid *→*Epi axes. Only miRNAs with direct CRC evidence or strong family-level EMT support are shown.

| miRNA | EMT Targets | Known CRC/EMT Role | Exosomal |
| --- | --- | --- | --- |
| miR-200a [22] | ZEB1/2, SMAD2/3, CTNNB1 | Canonical EMT suppressor, miR-200/ZEB loop | ✓<sup>a</sup> |
| miR-34a [23] | CTNNB1, SNAI1 | CRC EMT-associated suppressor via the IL-6R/STAT3/miR-34a axis | ✓<sup>b</sup> |
| miR-204 [24] | SNAI1/2, VIM, CDH1, MMP9 | CRC tumor suppressor; inhibits proliferation and invasion | — |
| miR-506 [25] | CTNNB1, SNAI1/2, VIM | CRC suppressor; inhibits proliferation, migration, and invasion | — |
| let-7g [26] | FN1, SMAD2, TGFBR1 | Tumor-suppressive let-7 family member; canonical RAS-regulating background | — |
| miR-708 [27] | MMP2, SMAD3, ZEB2 | CRC tumor suppressor; targets ZEB1 via AKT/mTOR signaling | — |
| miR-338 [28] | MMP9, ZEB2 | Tumor-suppressive / invasion-related miRNA in CRC | — |
| miR-129 [29] | CDH1, TWIST1, VIM | Linked to EMT regulation in CRC via the miR-129-5p/DNMT3A axis | — |

<sup>a</sup>Exosomal evidence is available for the miR-200 family in CRC (CAF-derived miR-200b-3p), rather than miR-200a specifically [30].
<sup>b</sup>MSC-derived extracellular vesicles carrying miR-34a-5p have been reported in CRC [31].

To rule out nonspecific enrichment, we performed Fisher’s exact tests comparing the 362 predicted targets of the top 10 miRNAs against seven curated pathway gene sets. Targets were most strongly enriched in EMT/plasticity (14/15 genes, OR = 793, *p* = 4.4 *×* 10^*−*24^), TGF-*β* signaling (7/12, OR = 78), Wnt/*β*-catenin (6/14, OR = 42), and PI3K-AKT (5/17, OR = 23), while genuinely unrelated negative control gene sets—ribosome, spliceosome, and oxidative phosphorylation—showed zero overlap (Fig. 4c). This result indicates that the predicted targeting is highly pathway-specific rather than a consequence of nonspecific enrichment among broadly expressed genes.

Finally, the target genes comprising this module are not uniformly expressed across epithelial cells but exhibit structured expression variation across Epi subclusters (Fig. 4a, right panel). Cluster C5, which has the highest plasticity score, shows elevated expression of mesenchymal-associated genes (FN1, VIM, TGFB1) and reduced expression of the epithelial marker CDH1, while other clusters display the inverse pattern. This indicates that mirCCC recovers not merely an abstract gene list, but a modular expression program with detectable heterogeneity in the receiver compartment. Notably, classical EMT transcription factors (SNAI1/2, ZEB1/2, TWIST1) are minimally expressed across all Epi subclusters, suggesting that this pattern is more consistent with a graded epithelial plasticity spectrum than with a complete canonical EMT program.

Together, these results support a receiver-centric communication program in which convergent stromal and myeloid miRNAs preferentially target an epithelial plasticity module in CRC epithelial cells.

## Discussion and conclusion

In this study, we developed mirCCC, a computational framework for inferring miRNA-mediated cell–cell communication from standard single-cell transcriptomic data. Unlike existing CCC methods that model protein ligand–receptor interactions, mirCCC reverse-estimates miRNA activity from target gene repression patterns and performs representation learning on per-miRNA directed communication graphs, incorporating the miRNA dimension into single-cell communication analysis.

A key feature of mirCCC is its repression-aware design, which ensures that high-scoring communication events depend on the concordance between sender proxy and receiver-side target gene repression, rather than on superficial expression correlations. In synthetic benchmarks, mirCCC was the only method whose performance improved upon the introduction of confounding cells, demonstrating robustness against false positives driven by surface-level sender/receiver characteristics. As community interest in miRNA communication inference grows, establishing standardized benchmark datasets will facilitate systematic comparison across methods.

Application to real data shows that mirCCC identifies biologically interpretable communication modules. In a CRC single-cell atlas [21], it recovered known CRC-associated miRNAs and revealed a receiver-centric program in which stromal and myeloid miRNAs converge on an EMT/TGF-*β*/Wnt/*β*-catenin-related plasticity module in epithelial cells, including IL-6R/STAT3/miR-34a signaling [32] and CAF-derived miR-200 exosomal signaling [30]. mirCCC identifies candidate miRNAs that converge between senders with pathway-specific targeting, illustrating its ability to generate modular hypotheses for downstream validation. Although based on a single MMRp patient, these results provide a foundation for assessing generalizability across multiple patients, cancer types, and non-tumor contexts. Minimal expression of canonical EMT transcription factors suggests mirCCC captures a graded epithelial plasticity spectrum rather than a full EMT program [2].

Several avenues for future research can be pursued. The current model treats all EV subtypes uniformly and does not distinguish exosomes from microvesicles [33, 34]. Integrating spatial transcriptomics could account for intercellular distance constraints [35, 36], and combining EV proteomics with small RNA-seq [37] would enable more comprehensive characterization of miRNA transfer mechanisms. Expanding the approach to multiple patients, cancer types, and cross-species scenarios could further enhance applicability. Overall, mirCCC provides a robust and interpretable framework, laying the groundwork for systematic studies of miRNA-mediated cell–cell communication.

## S1 Appendix. Supplementary benchmark methods and unified evaluation

Detailed descriptions of the unified prediction format, missing-pair completion procedure, scoring formulas, aggregation strategies, benchmark-specific adaptations, and parameter settings for mirCCC, Matched Product, Prior Product, miRTalk-Restored, CellChat-style, and NicheNet-style.

## S1 Table. Complete miRNA rankings across four analyzed communication axes

The table provides the complete miRNA rankings for the Strom-to-Epi, Myeloid-to-Epi, Epi-to-TNKILC, and Myeloid-to-TNKILC communication axes. For each miRNA, it reports the total communication score, rank, number of inferred cell-pair edges, mean score per edge, number of validated target genes detected in the dataset, and the number and identities of targets belonging to the curated epithelial-mesenchymal transition and epithelial plasticity gene set.

## Acknowledgments

We thank all the members of Ming Chen’s group for their valuable discussions. The authors have declared that no competing interests exist.

## Data availability

The colorectal cancer single-cell RNA-seq dataset used in this study (Pelka et al. 2021) is publicly available from the Gene Expression Omnibus under accession number GSE178341 (https://www.ncbi.nlm.nih.gov/geo/query/acc.cgi?acc=GSE178341). miRNA–target interaction data were obtained from miRTalk [12] (https://github.com/ZJUFanLab/miRTalk).

## Code availability

The code used to reproduce the experiments in this study is available at https://github.com/yfchen801/mirCCC. The repository includes the mirCCC analysis pipeline, benchmark evaluation code, synthetic data generation scripts, tutorial notebooks, and visualization utilities. Synthetic benchmark datasets can be reproduced using the scripts and configuration files provided in the repository.

## Author contributions

Conceptualization: Yifan Chen, Cong Feng, Ming Chen. Data Curation: Yifan Chen. Formal Analysis:Yifan Chen, Jiaming Cui. Funding Acquisition: Ming Chen. Investigation: Yifan Chen. Methodology: Yifan Chen, Jiaming Cui. Project Administration: Ming Chen, Cong Feng. Resources: Yifan Chen, Ming Chen, Shilong Zhang. Software:Yifan Chen, Jiaming Cui, Shilong Zhang. Supervision:Ming Chen, Cong Feng. Validation:Yifan Chen, Shilong Zhang. Visualization:Yifan Chen, Enyan Liu, Luyao Xie. Writing – Original Draft Preparation:Yifan Chen. Writing – Review & Editing:Ming Chen, Cong Feng, Yifan Chen.

